# An INSR-BECN1-KIAA0825 complex prevents hepatocytic insulin receptor secretion via extracellular vesicles against monogenic diabetes

**DOI:** 10.64898/2026.09.13.751259

**Authors:** Kenta Kuramoto, Min Chen, Congcong He

## Abstract

Monogenic diabetes from insulin receptor (INSR) mutations is a rare genetic disorder causing early-onset, profound insulin resistance. However, the pathogenic mechanisms by which INSR mutations disrupt receptor trafficking, cell-surface stability, and signaling are unresolved, and consequently, current treatment remains largely symptomatic and prognosis is not optimal. Here we discover that INSR, the autophagy protein BECN1, and KIAA0825, encoded by a diabetic-risk gene of previously unknown function, form a complex to regulate INSR transport, membrane presentation, and functionality in hepatocytes. Loss of hepatic BECN1 or KIAA0825, or expression of monogenic diabetes-causing mutations in INSR, disrupts the INSR-BECN1-KIAA0825 complex and mis-routes INSR to extracellular vesicles for secretion from hepatocytes, leading to impaired hepatocytic INSR cell-surface presentation and defects in hepatic insulin sensitivity, glycogen storage and exercise capacity. These findings identify new regulators and mechanisms governing INSR sorting, trafficking and signaling, uncover a non-canonical, non-degradative role for BECN1, and reveal the cellular fate of pathogenic INSR mutants underlying monogenic diabetes.

## Introduction

Different from type 1 or type 2 diabetes that result from abnormalities in multiple genes, monogenic diabetes is a rare genetic disorder caused by mutations in a single gene, for example, the insulin receptor (INSR) gene itself ^1^. INSR is a type II receptor tyrosine kinase that plays an essential role in glucose metabolism, development and growth ^2, 3^. In metabolic organs such as the liver, activation of INSR signaling is essential for stimulating glycogen synthesis from glucose up-taken from the bloodstream as nutrient and energy storage. Pathogenic mutations in INSR cause persistent hyperglycemia and insulin resistance of varying severity often manifesting in early childhood, leading to disorders including Type A insulin resistance syndrome (TAIRS), Donohue syndrome, and Rabson-Mendenhall syndrome ^1, 4, 5^. However, the cellular mechanisms by which INSR mutations impair INSR plasma-membrane presentation, receptor function, and signaling remain poorly defined, given INSR mRNA transcription and abundance are unaffected in the above disorders. In particular, how mutant INSR proteins are transported, regulated and retained in cells at the molecular level—and how these defective processes can be restored—are not well understood. As a result, current treatments for monogenic diabetes mainly include lifestyle modification (including diet and exercise) and management of symptoms and complications, but effective therapeutic cure to the disease is unavailable and the clinical prognosis remains poor ^1, 5^.

BECN1 is an autophagy protein and a core component of the autophagy-initiating class III phosphatidylinositol 3-kinase complex, which produces the lipid phosphatidylinositol (3)-phosphate to promote autophagosome formation. While autophagy proteins are generally known for their roles in lysosome-mediated cargo degradation, growing evidence suggests that they also participate in diverse cellular processes beyond this canonical pathway ^6–8^. Here, our study uncovers a novel, non-traditional mechanism for hepatocytic BECN1 in coordinating with KIAA0825, encoded by a diabetic-risk gene of previously unknown function, to regulate INSR cell-surface trafficking, stability and activity. INSR mutants identified in patients with monogenic diabetes evade the regulation of BECN1 and KIAA0825 and are diverted to extracellular vesicles/exosomes (EVs) for secretion, potentially to adjacent hepatic vascular endothelial cells (ECs). These findings reveal an unanticipated mechanism that regulates insulin receptor homeostasis and its disruption in monogenic diabetes.

## Results

### Liver BECN1 is essential for maintaining exercise capacity, hepatic insulin sensitivity, and glycogen storage

To study the metabolic function of hepatic BECN1, we generated WT control and liver-specific BECN1 knockout (KO) (BECN1^Δliver^) mice by intravenously injecting AAV8-TBG-Cre (Cre drive by the liver-specific TBG promoter) to WT or BECN1^flox/flox^ mice, respectively, validated by BECN1 protein depletion and accumulation of the autophagy cargo p62 (**Extended Data Fig. 1A**). We found that BECN1^Δliver^ mice show reduced maximal running capacity (**Fig. 1A**) and blood glucose level during exercise (**Fig. 1B**), compared to WT control mice. Intriguingly, in a fed-fasting-refed cycle after high-fat diet (HFD) challenge, although under fed conditions BECN1^Δliver^ mice were hyperglycemic compared to WT control mice, their blood glucose levels were significantly lower than that of WT mice during the fasting phase, despite comparable body weight at each timepoint as WT mice (**Fig. 1C**). However, such blood glucose decrease in BECN1^Δliver^ mice is not due to improved insulin sensitivity, because insulin-stimulated phosphorylation of Akt (leading to Akt activation) and glycogen synthase kinase-3β (leading to GSK3β inhibition) downstream of INSR is decreased, rather than increased, in the liver of BECN1^Δliver^ mice (**Fig. 1D**). Accordingly, we analyzed nutrient (carbohydrate and lipid) storage in these mice, and found that hepatic glycogen contents are reduced upon loss of hepatic BECN1 under all conditions tested, including fed and resting conditions, exercise, and fasting (**Fig. 1E**). In comparison, WT and BECN1^Δliver^ mice showed similar overall liver weight under either regular diet (RD) or HFD treatment (**Extended Data Fig. 1B-C**), and comparable hepatic triglyceride (TAG) contents (**Extended Data Fig. 1D**) synthesized from free fatty acids released from adipose tissue acutely induced by exercise or fasting as fuel ^9–11^, suggesting that hepatic BECN1 regulates carbohydrate, but not lipid, metabolism in response to exercise and fasting. It is also an unexpected finding that KO of BECN1 in the liver leads to a reduction, rather than an accumulation, of glycogen levels, given a role of autophagy in glycophagy, a form of selective glycogen degradation distinct from glycogenolysis ^12–14^. Thus, we reasoned that BECN1 regulates hepatic glycogen metabolism not through glycophagy, but via a previously unknown mechanism. We further found that although BECN1^Δliver^ mice have normal levels and distribution of GLUT2 (**Extended Data Fig. 1E-F**), the major liver glucose transporter ^15–17^, the mice showed markedly decreased hepatic glucose uptake (**Fig. 1F**). These data suggest that hepatic glycogen reduction in BECN1^Δliver^ mice is due to defective insulin-stimulated glycogen synthesis, thus leading to a failure in maintaining blood glucose and exercise capacity under stress conditions when breakdown of glycogen is needed, consistent with reported findings that liver glycogen levels positively correlate with exercise capacity ^9^. Thus, overall, these data demonstrate hepatic BECN1 as an important regulator of exercise capacity, liver insulin signaling and glycogen storage.

**Figure 1.**
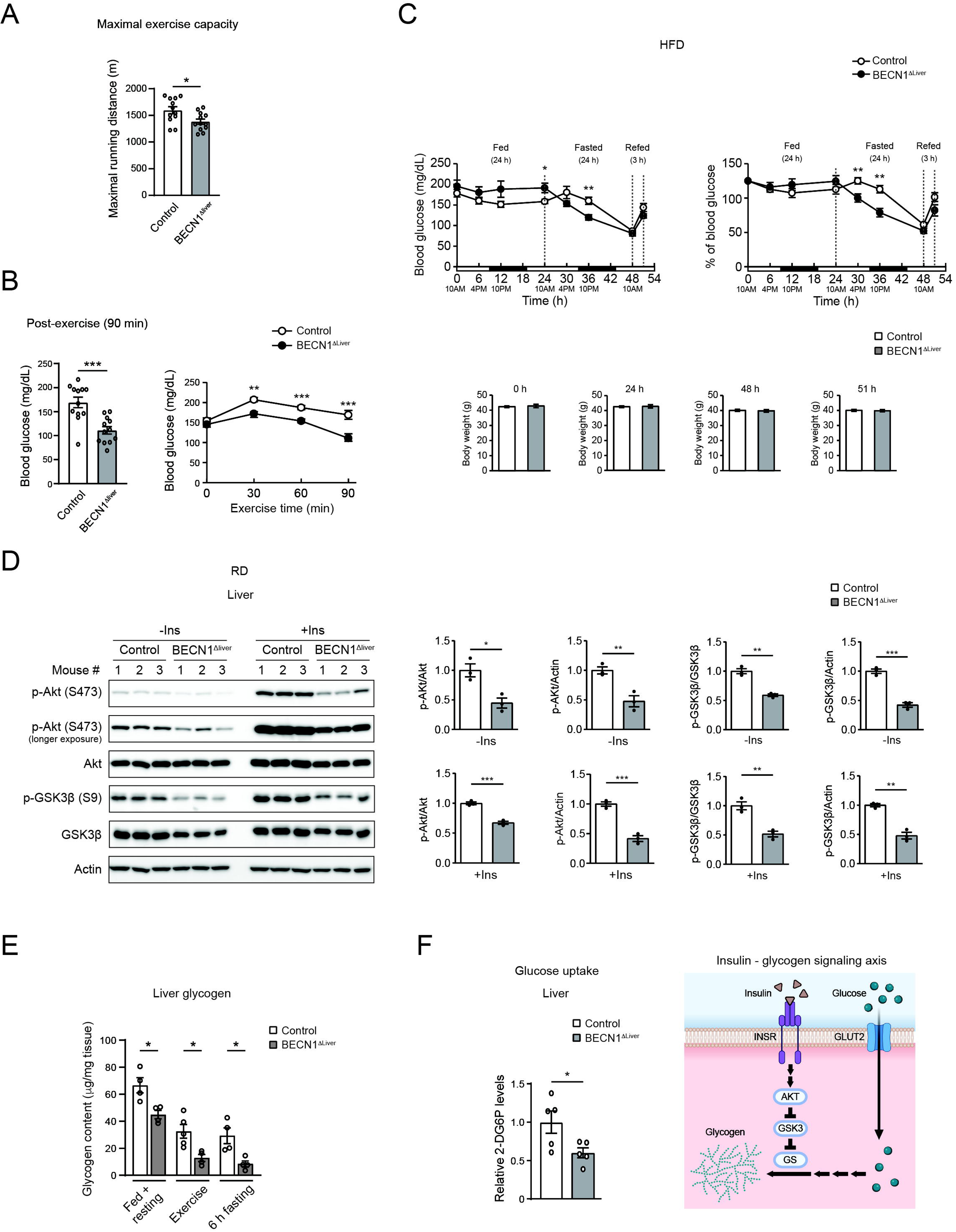
Loss of hepatic BECN1 reduces maximal exercise capacity, liver insulin sensitivity, and glycogen storage. **(A)** Reduced maximal running capacity of BECN1^Δliver^ mice compared to control mice under regular diet (RD)-fed conditions. N=12. T-test. **(B)** Blood glucose levels of RD-fed control and BECN1^Δliver^ mice during 90-min exercise. N=12. T-test. **(C)** Blood glucose levels and body weight of high-fat diet (HFD)-fed BECN1^Δliver^ mice and control mice during 24-h fed, 24-h fasting, and 3-h refed conditions. N=7-8. T-test. **(D)** Western blot analysis and quantification of insulin signaling (insulin-induced phosphorylation of Akt and GSK3β) in the liver of RD-fed control and BECN1^Δliver^ mice injected with insulin 15 min prior to tissue collection. -Ins, no insulin; +Ins, with insulin. **(E)** Glycogen storage levels in the liver of control and BECN1^Δliver^ mice under resting and fed conditions, after 90-min exercise, or after 6-h fasting. Two-way ANOVA with Tukey test. N=4-5. **(F)** Hepatic glucose uptake is reduced in BECN1^Δliver^ mice. 2-Deoxy-D-glucose (2-DG) levels were analyzed by metabolomics in the liver of control or BECN1^Δliver^ mice intraperitoneally co-injected with insulin, 2-Deoxy-D-glucose (2-DG), and glucose (to maintain physiological blood glucose) 30 min prior to tissue collection. Right, a diagram illustrating that insulin receptor (INSR) signaling increases hepatic glucose uptake and glycogen formation from glucose via Akt-induced phosphorylation and inhibition of GSK3 (Glycogen synthase kinase 3), the crucial negative regulator of the rate-limiting enzyme glycogen synthase (GS). Reduced insulin signaling leads to less glucose utilization and influx in the liver. N=5 mice. T-test. *, P<0.05; **, P<0.01; ***, P<0.001.

### BECN1 is essential for maintaining hepatic insulin receptor (INSR) cell-surface stability and signaling

To investigate the mechanism by which hepatic BECN1 regulates glycogen storage and insulin sensitivity, we screened for major players in a variety of metabolic pathways, and found that the level of mature form of insulin receptor (INSR, encoded by the INSR gene), but not other cytoplasmic metabolic regulators (including the mTOR pathway proteins, the circadian rhythm regulator CRY1, the ER stress marker GPR78/Bip, and endolysosomal markers EEA1 and LAMP1), is reduced in the liver of BECN1^Δliver^ mice (**Fig. 2A**, **Extended Data Fig. 2A**). Levels of additional cell-surface receptors, such as insulin-like growth factor 1 receptor (IGF1R) and epidermal growth factor receptor (EGFR), are not significantly altered by the loss of hepatic BECN1 (**Extended Data Fig. 2B**). Similarly, we observed a loss of mature INSR, but not IGF1R, in human Huh7 hepatic cells upon inducible depletion of BECN1 (**Fig. 2B**, **Extended Data Fig. 2C**). BECN1 knockdown (KD) also reduced INSR levels in another human hepatic cell line, HepG2 cells (**Extended Data Fig. 2D**). Confocal immunofluorescence imaging further demonstrated a loss of plasma membrane-localized INSR in BECN1^Δliver^ mouse liver (**Fig. 2C**). In addition, similar to RD-fed mice (**Fig. 1D**, **Extended Data Fig. 2E**), we found that BECN1 depletion also impairs insulin sensitivity (insulin-stimulated INSR and Akt phosphorylation) in the liver of HFD-fed mice (**Fig. 2D**), and in vitro in Huh7 and HepG2 hepatic cells (**Fig. 2E**, **Extended Data Fig. 2F**). Taken together, these data suggest that BECN1 is essential for the cell-surface presentation, stability, and function of the mature form of INSR in hepatocytes.

**Figure 2.**
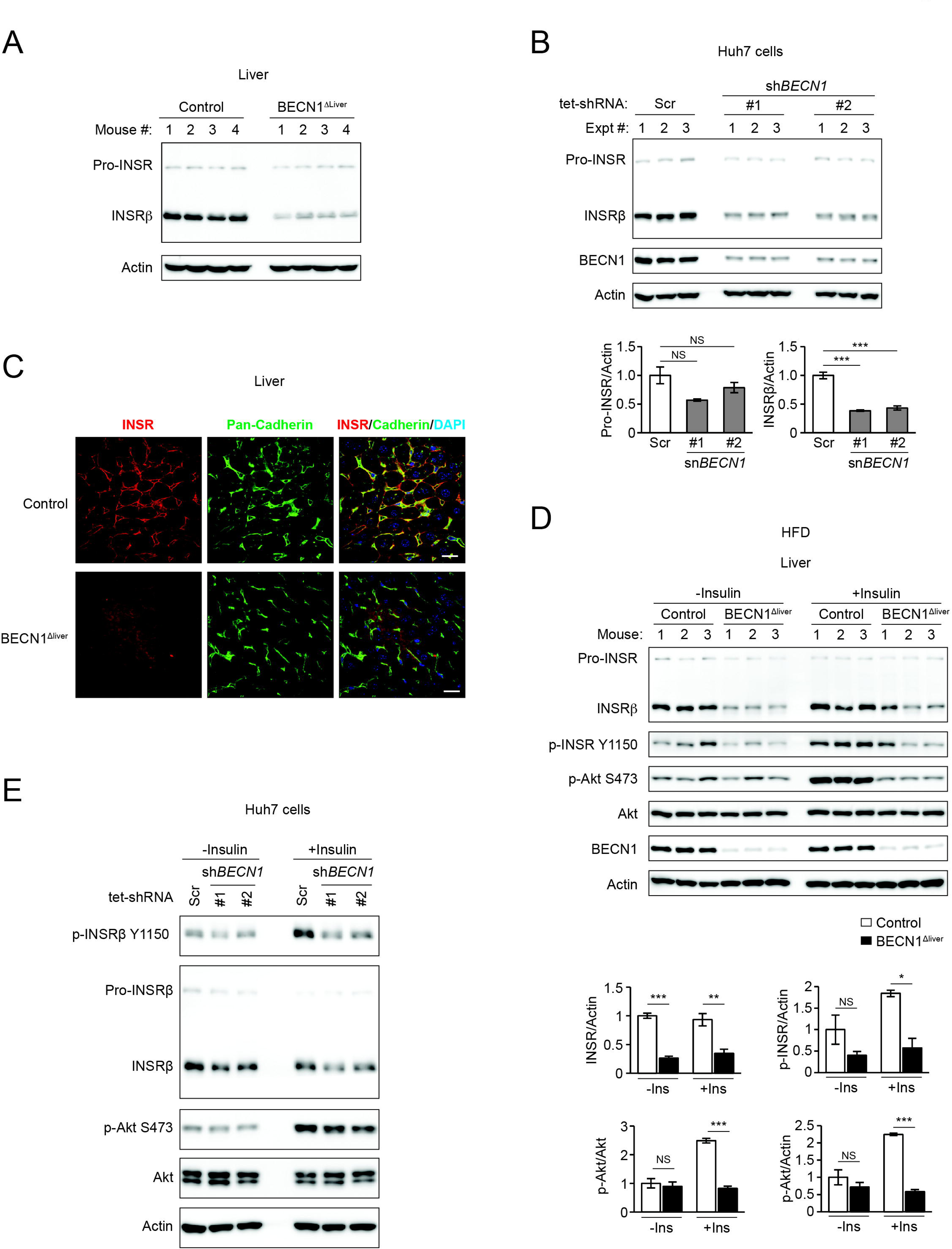
Loss of hepatic BECN1 reduces cell-surface insulin receptors (INSR) and insulin signaling. **(A)** Western blot analysis of pro- and mature INSR in the liver of BECN1^Δliver^ mice and control mice. **(B)** Western blot analysis and quantification of pro- and mature INSR in Huh7 cells stably expressing scrambled (Scr) or BECN1 shRNAs induced by 1 μg/ml tetracycline for 5 days. **(C)** Immunofluorescence imaging of INSR in the liver of RD-fed BECN1^Δliver^ mice and control mice. Cadherin staining illustrates the plasma membrane at cell-cell borders. Bar, 20 μm. **(D)** Western blot analysis and quantification of insulin-induced phosphorylation of INSR and Akt in the liver of BECN1^Δliver^ mice and control mice fed with 8 weeks of HFD and injected with 2 U/kg insulin 15 min prior to tissue collection. Two-way ANOVA with Tukey test. **(E)** Western blot analysis of insulin-induced phosphorylation of INSR and Akt in Huh7 cells stably expressing scrambled (Scr) or BECN1 shRNAs induced by 1 μg/ml tetracycline for 5 days, pre-sensitized with 0.5% FBS for 6 h, and treated with 100 nM insulin for 30 min prior to sample collection. *, P<0.05; **, P<0.01; ***, P<0.001.

### Hepatocytic BECN1 prevents mis-sorting of INSR to extracellular vesicles (EVs) for secretion from hepatocytes to liver vascular endothelial cells (ECs)

Next, we asked how BECN1 maintains INSR cell-surface presentation, as the finding is counterintuitive given BECN1 is an autophagy protein commonly considered to function in catabolism. BECN1 depletion does not affect INSR mRNA transcription, both in vivo (BECN1^Δliver^ mouse liver) (**Extended Data Fig. 3A**) and in vitro (Huh7 hepatic cells) (**Extended Data Fig. 3B**). To explore whether INSR can be secreted out of the cells, we isolated extracellular vesicles (EVs) from conditioned medium of control and BECN1-KD Huh7 cells. We found that loss of BECN1 increases EV-incorporation of both endogenous (**Fig. 3A**) and transfected (**Extended Data Fig. 3C**) INSR. EVs purified from the plasma of BECN1^Δliver^ mice (**Fig. 3B**), or conditioned medium of BECN1-KD HepG2 cells (**Extended Data Fig. 3D**), also contained elevated levels of mature INSR. KD of RAB27A, a small GTPase required for EV secretion ^18^, reduces EV incorporation of INSR induced by BECN1 depletion (lane 2 vs. lane 4, **Extended Data Fig. 3E**), supporting that BECN1 functions to prevent RAB27A-facilitated EV recruitment of INSR. Furthermore, we found that C2C12 myotubes cultured with EVs purified from BECN1^Δliver^ mouse plasma have an elevated INSR protein level, compared to those cultured with EVs from plasma of WT mice (**Fig. 3C**), suggesting that as recipient cells, C2C12 myotubes received more INSR delivered by EVs from BECN1^Δliver^ mice than from WT mice. Importantly, the EV-delivered INSR is functional, evidenced by increased insulin signaling (INSR phosphorylation) in the myotubes cultured with BECN1^Δliver^ mouse plasma EVs (**Fig. 3C**). Consistently, EVs isolated from conditioned medium of BECN1-KD Huh7 cells also increase INSR levels in recipient C2C12 myotubes, compared to EVs secreted from WT Huh7 cells (**Fig. 3D**). Thus, these data suggest that BECN1 depletion leads to misrouting and secretion of functional INSR in EVs.

**Figure 3.**
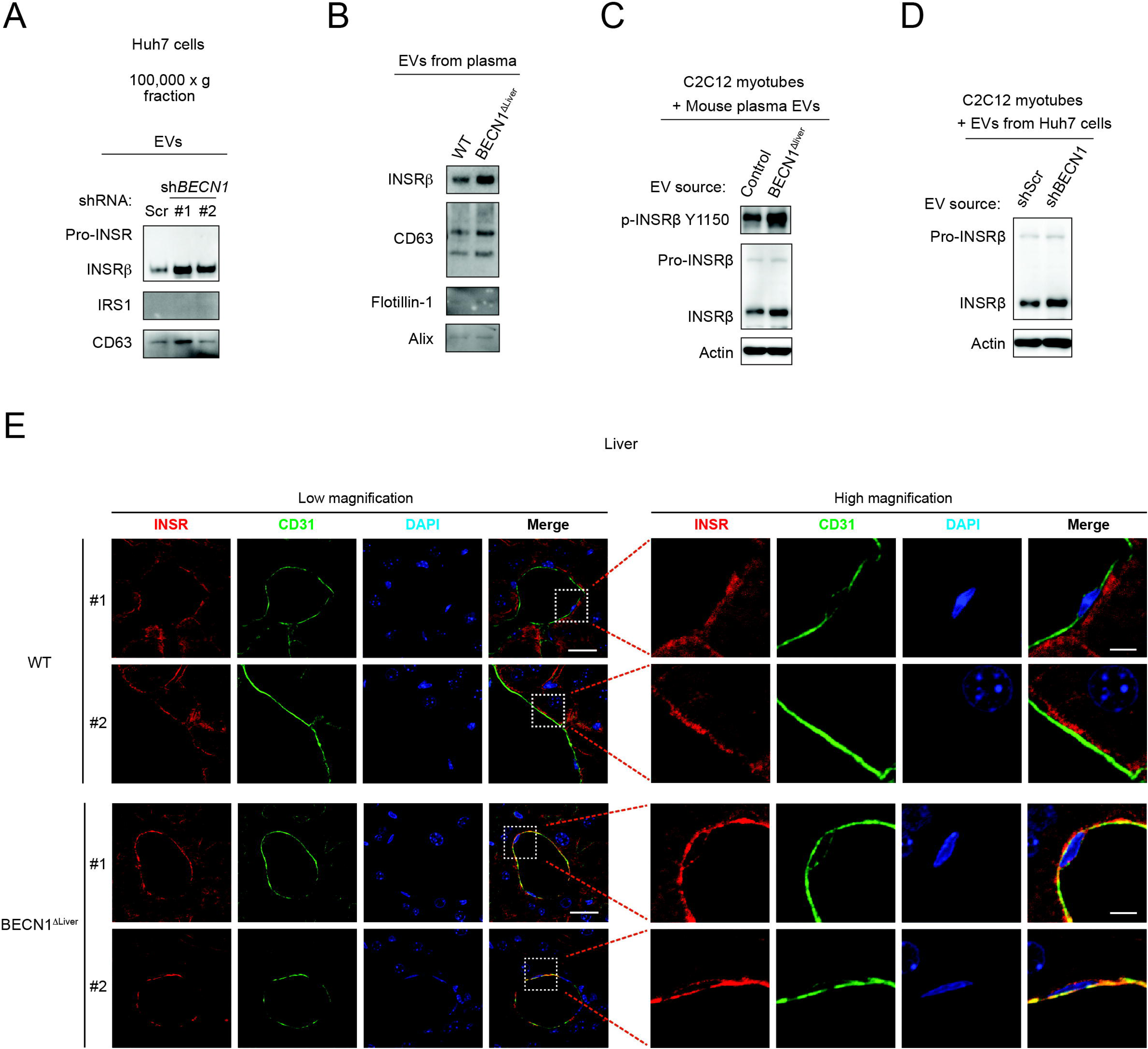
Hepatocyte-specific BECN1 KO re-routes INSR to EVs, leading to reduced INSR in hepatocytes and elevated INSR in liver vascular endothelial cells. **(A)** Western blot analysis of pro- and mature INSR in EVs isolated from conditioned media of Huh7 cells expressing scrambled (Scr) or two different BECN1 shRNAs. **(B)** Western blot analysis of INSR in EVs isolated from plasma of RD-fed BECN1^Δliver^ mice and control mice. CD63, Flotillin-1 and Alix serve as EV markers. **(C)** Western blot analysis of INSR levels and activation (insulin-stimulated INSR phosphorylation) in WT C2C12 myotubes cultured with EVs isolated from the plasma of BECN1^Δliver^ mice or control mice for 16 h. **(D)** Western blot analysis of INSR in WT C2C12 myotubes cultured for 16 h with EVs isolated from conditioned media of Huh7 cells stably expressing scrambled (Scr) or BECN1 shRNA. **(E)** Immunofluorescence confocal imaging of INSR and CD31 (endothelial marker) in the liver of BECN1^Δliver^ mice and control mice. Low magnification bar, 20 μm; high magnification bar, 5 μm.

An intriguing question is the destination of hepatocyte-released EVs in the absence of BECN1 in vivo. To address this question, we performed confocal imaging of INSR in various cell types in the liver and additional metabolic tissues of control vs. BECN1^Δliver^ mice. We found that different from control mice, where hepatocytes show the strongest cell-surface INSR signals in the liver (INSR and vascular EC marker CD31 signals are adjacent but not colocalized/overlapped, **Fig. 3E** upper panel), in BECN1^Δliver^ mouse liver, the cell-surface INSR signal is lost in hepatocytes, but strongly presented in adjacent vascular ECs (INSR and CD31 now display substantial colocalization, **Fig. 3E** lower panel). Thus, hepatocyte-specific BECN1 KO reduces INSR in hepatocytes while elevating cell-surface INSR in hepatic vascular ECs, consistent with enhanced EV-mediated transfer of INSR from hepatocytes to hepatic ECs. In comparison, KO of BECN1 in hepatocytes does not significantly increase cell-surface INSR levels in vascular ECs of more distant organs, such as the brown adipose tissue (BAT) (**Extended Data Fig. 3F**), suggesting that the non-cell autonomous effects of BECN1 KO on INSR plasma-membrane presentation are localized or short-distance. We reasoned that hepatocyte-derived INSR-containing EVs are effectively received by hepatic vascular ECs before reaching remote organs, thus demonstrating a paracrine patten.

### An INSR-BECN1-KIAA0825 complex stabilizes INSR at the plasma membrane and prevents its recruitment and secretion by EVs

To determine the regulatory mechanism of BECN1 in preventing INSR secretion, we performed confocal imaging for potential colocalization between INSR and BECN1 in hepatic HepG2 cells. We found that in addition to INSR’s predominant plasma membrane localization, INSR also localizes to a number of cytoplasmic puncta, which colocalize with BECN1 (tagged with either N-terminal or C-terminal RFP) (**Fig. 4A**, **Extended Data Fig. 4A**). These data suggest that BECN1 and INSR may interact in hepatocytes. Via co-immunoprecipitation (co-IP) assays, we found that BECN1 is pulled down by INSR in both cell lysates (**Fig. 4B**) and membrane fractions (**Fig. 4C**) of Huh7 cells, supporting that INSR and BECN1 form a complex. Co-IP of INSR and BECN1 was also detected in cell lysates and membrane factions when IgG (instead of empty vector) was used as negative control (**Extended Data Fig. 4B-C**). We also detected the interaction in vivo between endogenous BECN1 and INSR in the mouse liver via co-IP (**Fig. 4D**). Together, these data suggest that INSR interacts with BECN1 both in vitro and in vivo.

**Figure 4.**
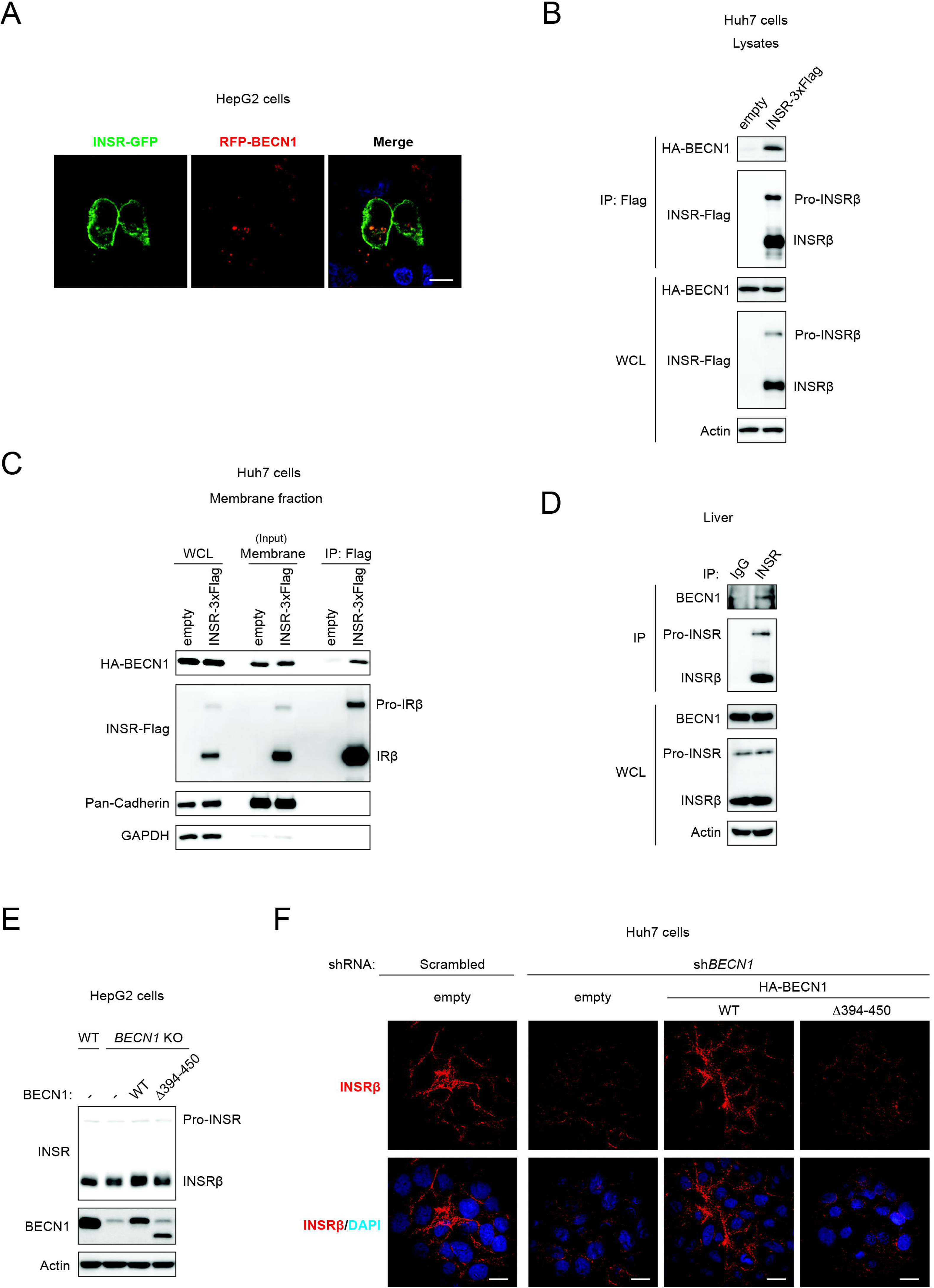
INSR interacts with BECN1, which is essential for INSR intracellular levels and plasma-membrane presentation. **(A)** Confocal fluorescence microscopy of INSR and BECN1 in HepG2 cells showing cytosolic punctum colocalization between INSR-GFP and RFP-BECN1 (N-terminal RFP tag). Bar, 10 μm. **(B)** Co-immunoprecipitation (co-IP) of BECN1 with INSR in lysates of Huh7 cells transfected with HA-BECN1 and INSR-Flag. **(C)** Co-IP of BECN1 with INSR in the membrane fractions of Huh7 cells transfected with HA-BECN1 and INSR-Flag. Empty vector (no INSR-Flag) transfection serves as a negative control. **(D)** Co-IP of endogenous BECN1 with endogenous INSR in the liver of WT mice. **(E)** Western blot analysis of INSR levels in WT HepG2 cells, or BECN1 KO HepG2 cells transfected with empty vector, WT BECN1, or BECN1^Δ394–450^. **(F)** Immunofluorescence imaging of INSR in Huh7 cells stably expressing scrambled shRNA transfected with empty vector, or stably expressing BECN1 shRNA transfected with empty vector, WT BECN1, or BECN1^Δ394–450^. Bar, 20 μm. WCL, whole cell lysates.

To reveal the physiological importance of INSR-BECN1 binding, we performed interaction domain mapping by co-IP, using truncation and deletion mutants of BECN1, and found that the C-terminal 394-450 amino acid region of BECN1 is required for binding INSR (**Extended Data Fig. 4D**). In contrast to WT BECN1, which rescued both the protein level (**Fig. 4E**) and cell-surface localization (**Fig. 4F**, **Extended Data Fig. 4E**) of INSR in BECN1 KO/KD HepG2 or Huh7 cells, the loss-of-INSR interaction BECN1^Δ394–450^ mutant failed to do so. These data suggest that BECN1-INSR interaction is essential for INSR stability and plasma-membrane presentation.

We further found that the INSR-BECN1 interaction is dynamic and can be enhanced by conditions such as serum deprivation (**Extended Data Fig. 5A**), suggesting that additional players exist to regulate the INSR-BECN1 complex. To fully characterize the INSR-BECN1 complex, we performed tandem IP for both BECN1-HA and INSR-Flag in Huh7 cells followed by mass spectrometry analysis, in order to identify new partners that simultaneously bind both BECN1 and INSR (**Extended Data Fig. 5B**). We discovered KIAA0825, encoded by a gene of previously unknown function that is associated with autosomal recessive postaxial polydactyly ^19–22^ and reported as a diabetes-risk gene in genome-wide association studies (GWASs) of diverse human populations ^23, 24^. We discovered that the KIAA0825 gene encodes two protein isoforms, the long isoform KIAA0825(L) (∼150 kDa with 1297 amino acids) and the short isoform KIAA0825(S) (∼38 kDa with 324 amino acids), both of which express in the mouse liver (**Extended Data Fig. 5C**). By co-IP, we validated that both KIAA0825 isoforms coimmunoprecipitate with either BECN1 (**Fig. 5A**, **Extended Data Fig. 5D**), or INSR (**Fig. 5B**, **Extended Data Fig. 5E**), or both BECN1 and INSR (**Fig. 5C**, **Extended Data Fig. 5F**) in Huh7 cell lysates. The KIAA0825-BECN1 interaction is also detected in the membrane factions of Huh7 cells (**Fig. 5D**). Depletion of KIAA0825 (both long and short isoforms) via shRNAs (**Extended Data Fig. 5G**) reduces INSR levels and INSR-BECN1 binding in cell lysates (WCL) of Huh7 cells, and simultaneously increases INSR levels in the EVs purified from the conditioned medium (**Fig. 5E**, **Extended Data Fig. 5H**). In comparison, overexpression of KIAA0825 had opposite effects on INSR levels in the EVs (**Fig. 5F**). In line with these findings, confocal microscopy further revealed that KIAA0825 KO drastically decreases the cell-surface localization of INSR in hepatic HepG2 cells (**Fig. 5G**). Overall, these data suggest that KIAA0825 plays an important role in maintaining cell-surface INSR and preventing INSR recruitment and secretion by EVs. Altogether, we conclude that the INSR-BECN1-KIAA0825 complex is an essential molecular controller for INSR transport and stabilization at the plasma membrane, but not to EVs.

**Figure 5.**
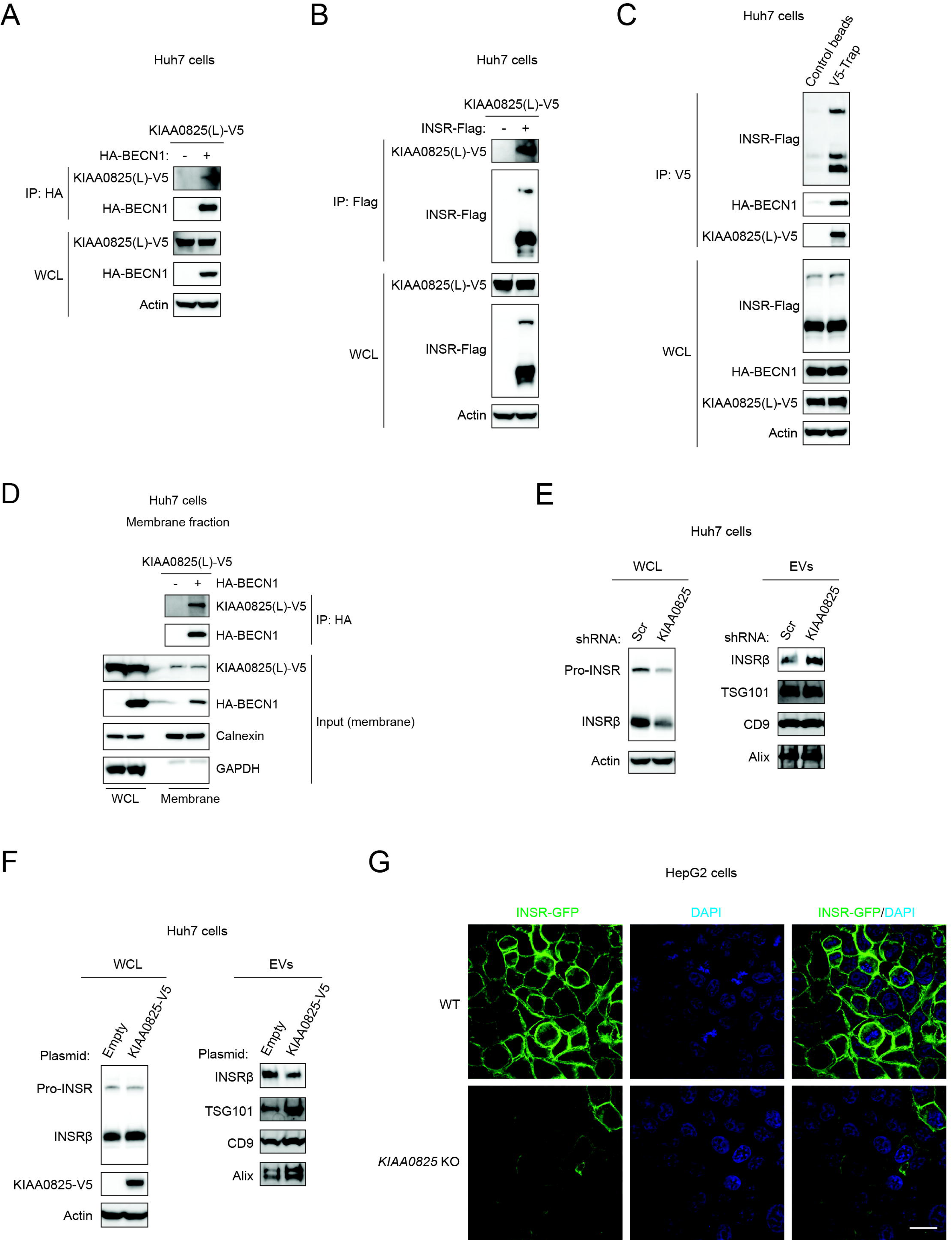
KIAA0825, encoded by a diabetic risk gene of unknown function, is a component of the INSR-BECN1 complex, and prevents EV incorporation and secretion of INSR. **(A)** Co-IP of KIAA0825(L) (long isoform) by BECN1 in Huh7 cells transfected with HA-BECN1 and KIAA0825(L)-V5. **(B)** Co-IP of KIAA0825(L) by INSR in Huh7 cells transfected with INSR-Flag and KIAA0825(L)-V5. **(C)** Co-IP of INSR and BECN1 by KIAA0825(L) in Huh7 cells transfected with INSR-Flag, HA-BECN1 and KIAA0825(L)-V5. **(D)** Co-IP of KIAA0825(L) by BECN1 in the membrane fractions of Huh7 cells transfected with KIAA0825(L)-V5 and HA-BECN1. **(E)** Western blot analysis of INSR in WCL (whole cell lysates) or EVs isolated from conditioned media of Huh7 cells stably expressing scrambled (Scr) or KIAA0825 shRNA. **(F)** Western blot analysis of INSR in WCL (whole cell lysates) or EVs isolated from conditioned media of Huh7 cells overexpressing an empty vector or KIAA0825 protein. **(G)** Confocal imaging of INSR in WT and KIAA0825 KO HepG2 cells stably expressing INSR-GFP. Bar, 20 μm.

### Rare monogenic diabetic INSR mutations disrupt the INSR-BECN1-KIAA0825 complex, cause EV incorporation and secretion of INSR, and reduce INSR cell-surface availability

Rare mutations in the subunit β of INSR, such as R1020Q, A1055V and R1191Q, lead to insulin resistance and monogenic diabetes in humans (https://web.expasy.org/variant_pages/VAR_004092.html, https://web.expasy.org/variant_pages/VAR_015923.html, https://web.expasy.org/variant_pages/VAR_004098.html) ^25–27^. These monogenic diabetic mutations are located in the intracellular tyrosine kinase domain (TK) and the juxtamembrane (JM) domain that connects the transmembrane domain (TM) to TK (**Fig. 6A**). However, how these INSR mutations cause diseases, and how they are molecularly regulated, are poorly understood. Via co-IP analyses, we found that compared to WT INSR, all three INSR mutants tested showed a reduced interaction with both BECN1 (**Fig. 6B**, **Extended Data Fig. 6**) and KIAA0825 (**Fig. 6B**), suggesting that these diabetic INSR mutations disrupt the INSR-BECN1-KIAA0825 complex. Instead of cell-surface presentation as WT INSR, the R1020Q, A1055V and R1191Q mutants of INSR colocalized with CD63, a marker for EVs that are formed in multivesicular bodies (MVBs), suggesting that the INSR mutants are mis-sorted to EVs upon loss of BECN1 and KIAA0825 interactions (**Fig. 6C**). As a result, we found that compared to WT INSR, R1020Q, A1055V and R1191Q mutations in INSR cause increased incorporation of INSR into EVs purified from the conditioned medium of hepatic Huh7 cells (**Fig. 6D**). Thus, these data suggest that the rare INSR mutants identified in patients with monogenic diabetes evade the BECN1-KIAA0825 regulation, are misrouted into EVs for secretion, and have impaired plasma-membrane presentation.

**Figure 6.**
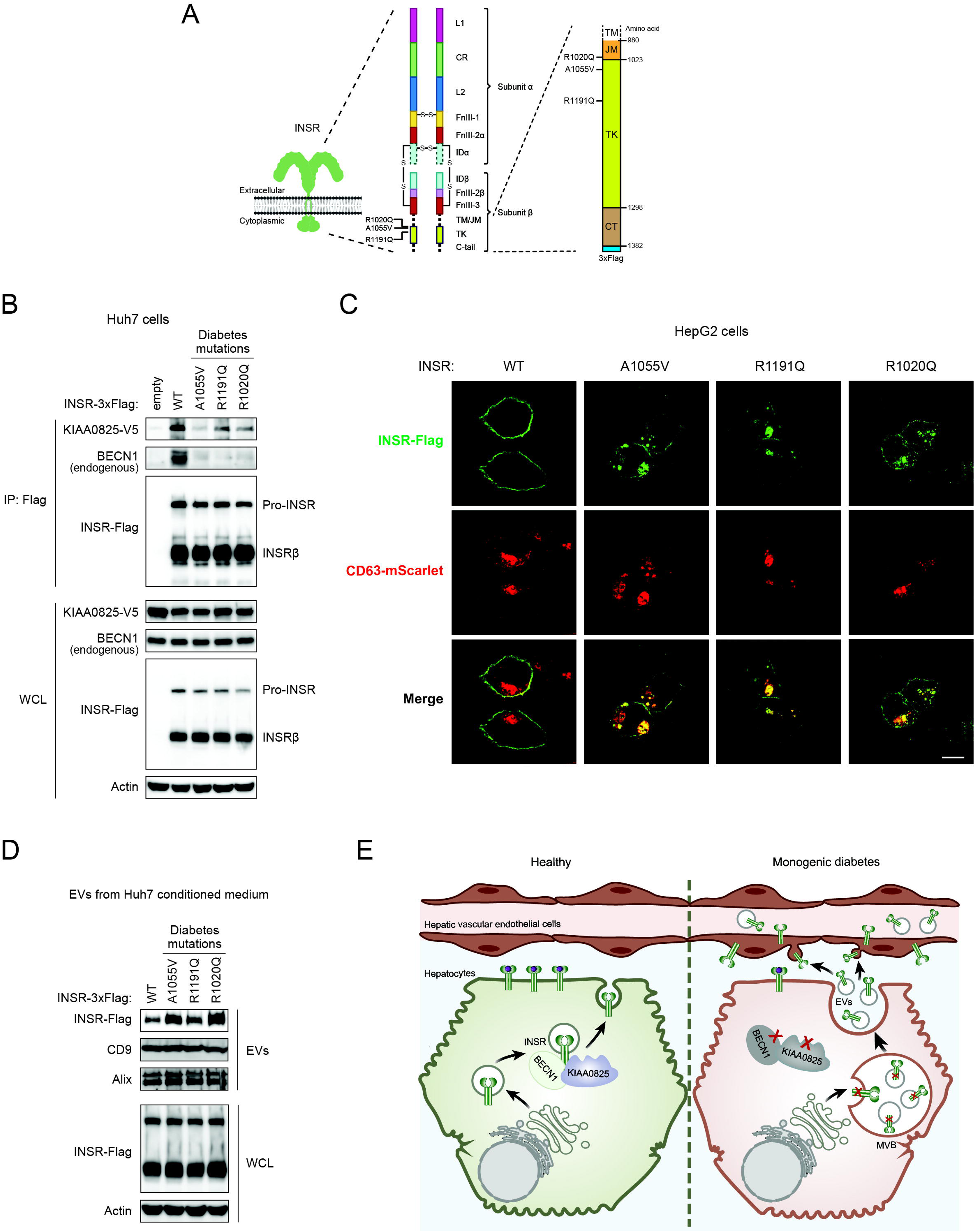
Monogenic diabetic INSR mutations disrupt the INSR-BECN1-KIAA0825 complex, misroute INSR to EV recruitment and secretion, and reduce INSR plasma-membrane presentation. **(A)** Schematic model of INSR and locations of rare INSR mutations (R1020Q, A1055V, R1191Q) identified in patients of monogenic diabetes. INSR is a heterotetramer of 2 subunits α and 2 subunits β, where subunit α binds the ligand insulin and subunit β mediates kinase signaling. The INSR gene encodes immature pro-INSR, synthesized in the early secretory pathway (ER/Golgi). Pro-INSR is further cleaved in the trans-Golgi network by furin or a related subtilisin-like proprotein convertase into subunit α and subunit β, which are covalently linked by 4 disulfide bonds (-S-S-) to form mature INSR. TM/JM: trans-and juxta-membrane domains; TK: tyrosine kinase domain; C-tail: C-terminal tail; L1, L2: 1^st^ and 2^nd^ leucine-rich-repeat domains; CR: cysteine-rich domain; FnIII-1, -2, -3: 1^st^, 2^nd^ and 3^rd^ type III fibronectin domains; IDα and IDβ: α-chain and β-chain components of the insert domain. **(B)** Co-IP of KIAA0825 and endogenous BECN1 with human WT INSR or INSR mutants (A1055V, R1191Q and R1020Q identified in monogenic diabetic patients) in Huh7 cells transfected with KIAA0825-V5 and INSR-Flag. **(C)** Immunofluorescence microscopy of INSR cell-surface and intracellular localization in HepG2 cells expressing WT INSR or diabetic INSR mutants (A1055V, R1191Q and R1020Q). CD63 serves as an MVB/EV marker. Bar, 10 μm. **(D)** Western blot analysis of INSR levels in EVs isolated from conditioned media of Huh7 cells expressing WT INSR or diabetic INSR mutants (A1055V, R1191Q and R1020Q). CD9 and Alix serve as EV markers. **(E)** Schematic model of INSR regulation by BECN1 and KIAA0825. BECN1, KIAA0825, and INSR form a complex to promote cell-surface transport and stability of INSR in normal hepatocytes (left). Loss of BECN1 or KIAA0825, or expression of diabetes-causing mutations in the INSR cytosolic region, disrupt the INSR-BECN1-KIAA0825 complex and divert INSR to EVs (Right), which can be absorbed by hepatic vascular endothelial cells. Such mis-sorting leads to reduced hepatocytic INSR and increased endothelial INSR, hepatic insulin resistance, and defects in glycogen storage and exercise capacity. MVBs, multivesicular bodies.

## Extended Data Figure Legends

**Extended Data Figure 1.**
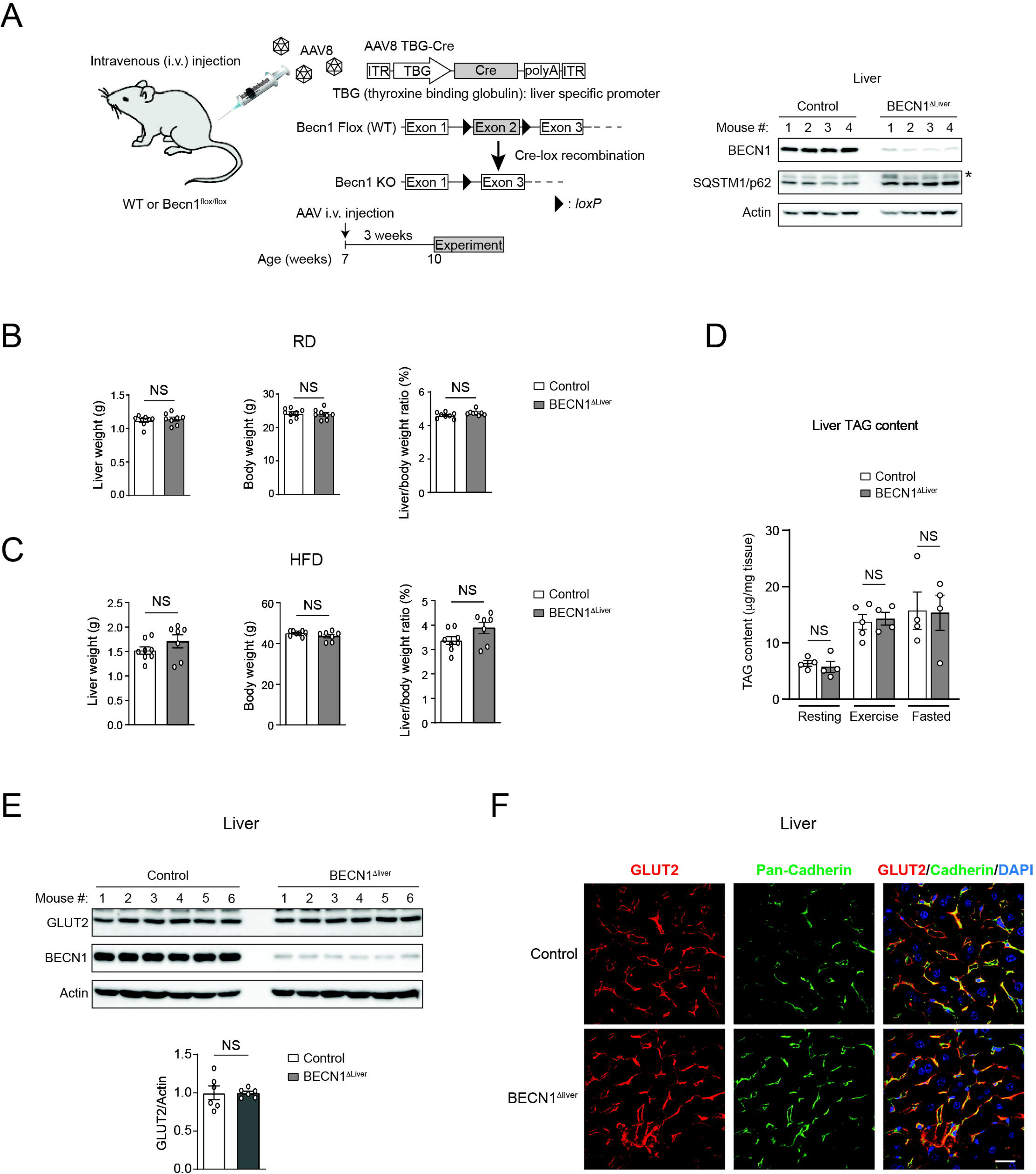
Characterization of BECN1^Δ^^liver^ mice. **(A)** Generation of BECN1^Δliver^ mice. AAV8-TBG-Cre was intravenously (i.v.) injected into 7-week old WT or BECN1^flox/flox^ mice, resulting in WT control and BECN1^Δliver^ mice, respectively. Experiments were performed 3-7 weeks after AAV injection. Hepatic depletion of BECN1 and accumulation of p62 were validated by Western blot analysis. *, non-specific band. **(B)** Liver weight and liver/body weight percentage are comparable in RD-fed control and BECN1^Δliver^ mice. N=8. T-test. **(C)** Liver weight and liver/body weight percentage are comparable in control and BECN1^Δliver^ mice fed with 12-week HFD. N=7-8. T-test. **(D)** Liver TAG (triglyceride) contents are comparable in RD-fed control and BECN1^Δliver^ mice. Two-way ANOVA with Tukey test. N=4-5. **(E)** Western blot analyses showing comparable levels of GLUT2, the major liver glucose transporter and a passive transporter, in RD-fed control and BECN1^Δliver^ mice. N=6. T-test. **(F)** Immunofluorescence imaging of GLUT2 in liver sections of RD-fed control and BECN1^Δliver^ mice. Cadherin staining illustrates the plasma membrane at cell-cell borders. Bar, 20 μm.

**Extended Data Figure 2.**
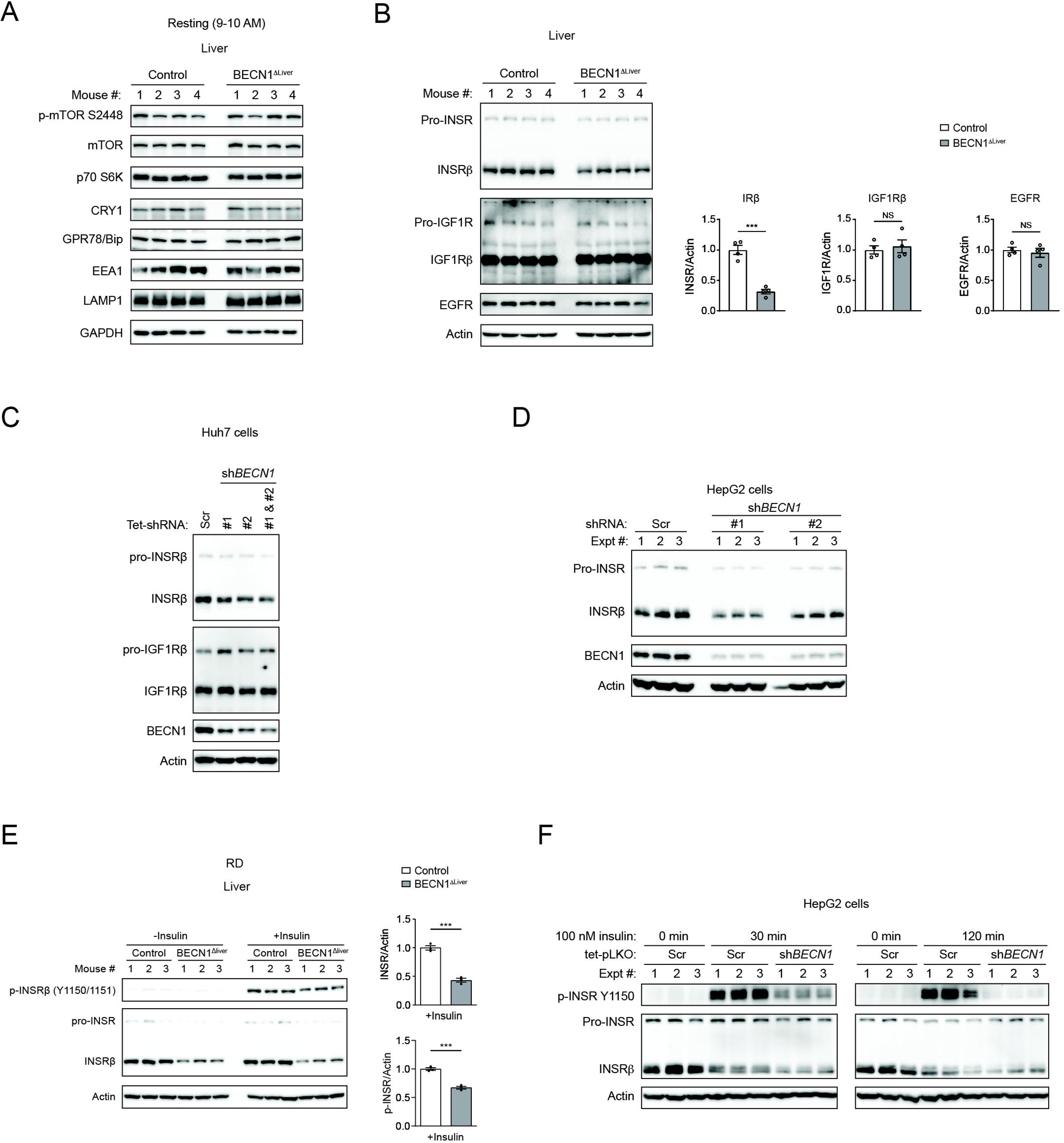
Loss of BECN1 decreases INSR levels and sensitivity in hepatic cells and in vivo. **(A)** Western blot analysis of multiple cellular markers (including the mTOR, circadian rhythm, ER stress, and endolysosomal pathways) in the liver of RD-fed control and BECN1^Δliver^ mice. **(B)** Western blot analysis and quantification of pro- and mature INSR, pro- and mature IGF1R, and EGFR in the liver of RD-fed control and BECN1^Δliver^ mice. N=4. T-test. **(C)** Western blot analysis of INSR and IGF1R in Huh7 cells stably expressing tetracycline (Tet)-inducible scrambled (Scr) shRNA or two different BECN1 shRNAs (1 μg/ml tetracycline treatment for 5 days). **(D)** Western blot analysis of pro- and mature INSR in HepG2 cells stably expressing scrambled or two different BECN1 shRNAs. **(E)** Western blot analysis of INSR levels and activation in HepG2 cells stably expressing Tet-inducible scrambled or BECN1 shRNA, stimulated with 100 nM insulin for 0 min, 30 min and 120 min. **(F)** Western blot analysis and quantification of INSR levels and activation (p-INSR) in the liver of RD-fed control and BECN1^Δliver^ mice injected with insulin 15 min prior to tissue collection. ***, P<0.001; NS, not significant.

**Extended Data Figure 3.**
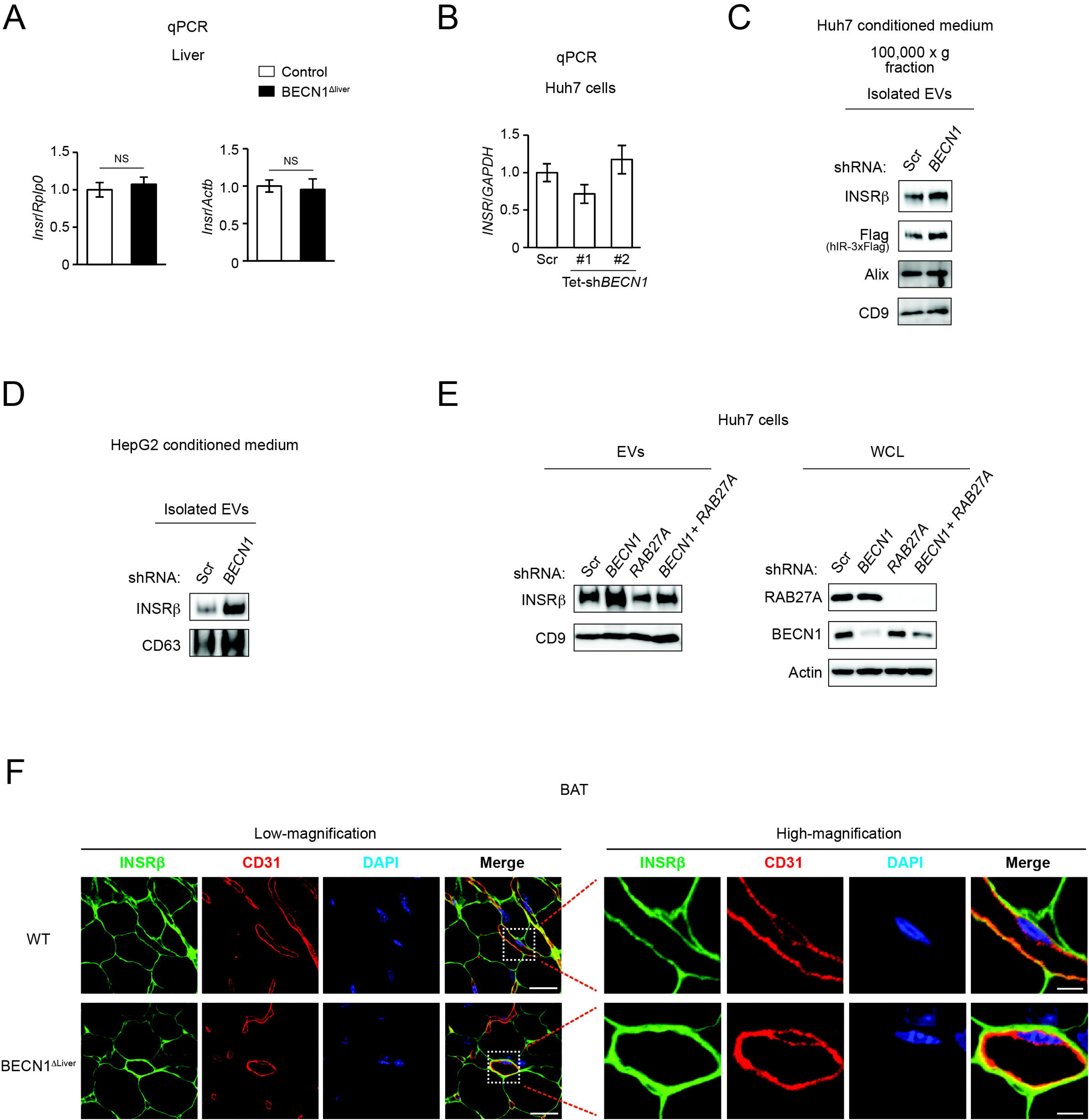
Loss of BECN1 promotes INSR translocation into the EVs. **(A)** qPCR analysis of INSR transcriptional levels in the liver of RD-fed control and BECN1^Δliver^ mice. N=4. T-test. **(B)** qPCR analysis of INSR transcriptional levels in Huh7 cells stably expressing Tet-inducible scrambled (Scr) shRNA or two different BECN1 shRNAs (1 μg/ml tetracycline treatment for 5 days). N=6. T-test. **(C)** Western blot analysis of INSR in EVs isolated from conditioned media of Huh7 cells stably expressing scrambled or BECN1 shRNA and transiently transfected with INSR-Flag. **(D)** Western blot analysis of endogenous INSR in EVs isolated from conditioned media of HepG2 cells stably expressing scrambled or BECN1 shRNA. **(E)** Western blot analysis of INSR levels in EVs isolated from conditioned media of Huh7 cells stably expressing scrambled, BECN1, RAB27A, or both BECN1 and RAB27A (double knockdown) shRNAs. Gene knockdown efficiency was analyzed by Western blot studies in WCL (whole cell lysates). **(F)** Immunofluorescence confocal imaging of INSR and CD31 (endothelial marker) in the brown adipose tissue (BAT) of BECN1^Δliver^ mice and control mice. Low magnification bar, 20 μm; high magnification bar, 5 μm.

**Extended Data Figure 4.**
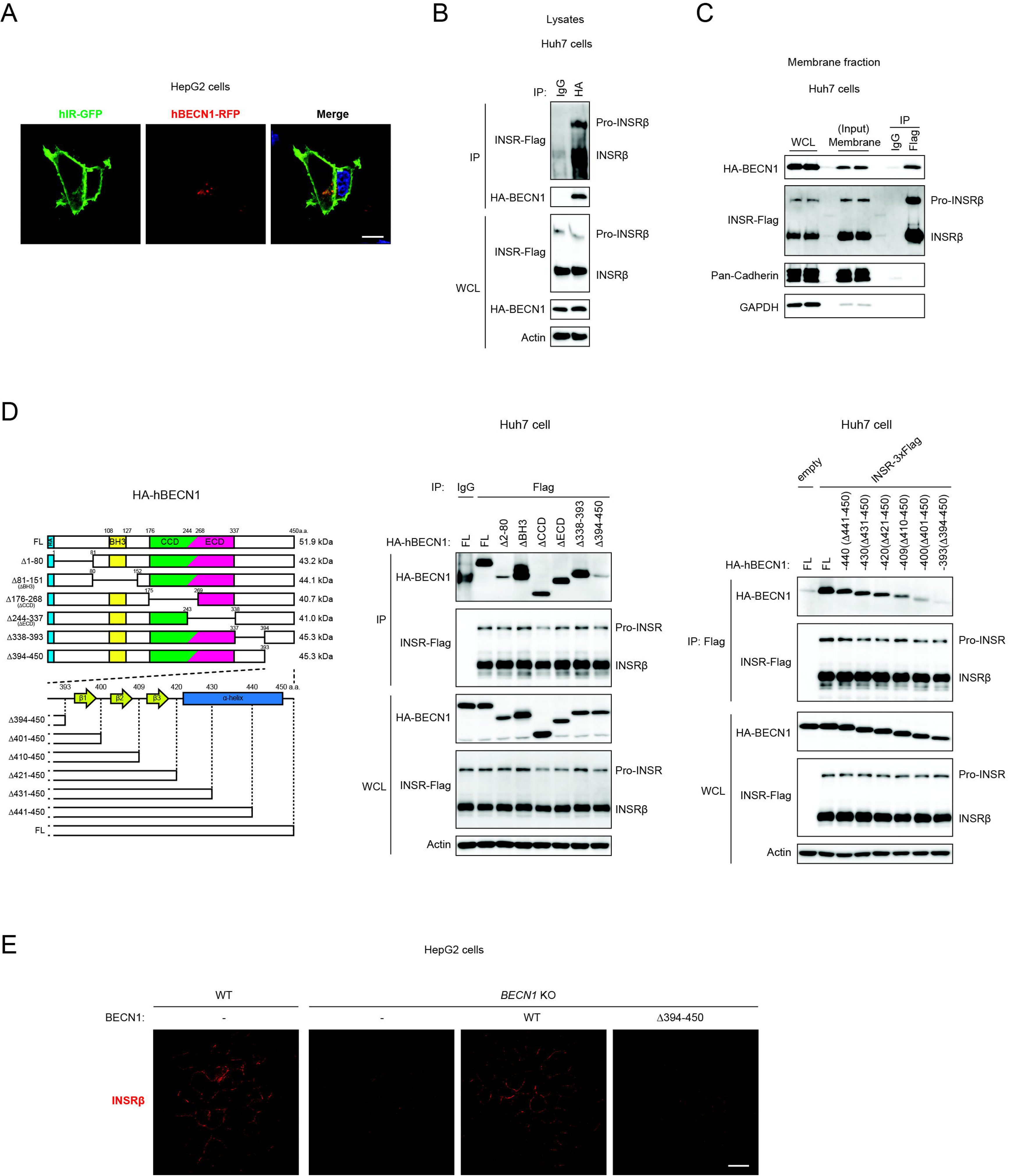
INSR interacts with BECN1, which is essential for the cell-surface presentation of INSR. **(A)** Confocal fluorescence microscopy of INSR and BECN1 in HepG2 cells showing cytosolic punctum colocalization between INSR-GFP and BECN1-RFP (C-terminal RFP tag). Bar, 10 μm. **(B)** Co-IP of INSR by BECN1 in lysates of Huh7 cells transfected with HA-BECN1 and INSR-Flag. IgG IP serves as a negative control. **(C)** Co-IP of BECN1 by INSR in the membrane fractions of Huh7 cells transfected with HA-BECN1 and INSR-Flag. IgG IP serves as a negative control. **(D)** Co-IP of HA-tagged full-length (FL) or indicated deletion/truncation mutants of BECN1 with Flag-tagged INSR in Huh7 cells. Left, IgG IP serves as a negative control. Right, empty vector (without INSR-Flag) transfection serves as a negative control. **(E)** Immunofluorescence microscopy of INSR in WT HepG2 cells, and BECN1 KO HepG2 cells transfected with empty vector, WT BECN1 or the BECN1^Δ394–450^ mutant. Bar, 20 μm.

**Extended Data Figure 5.**
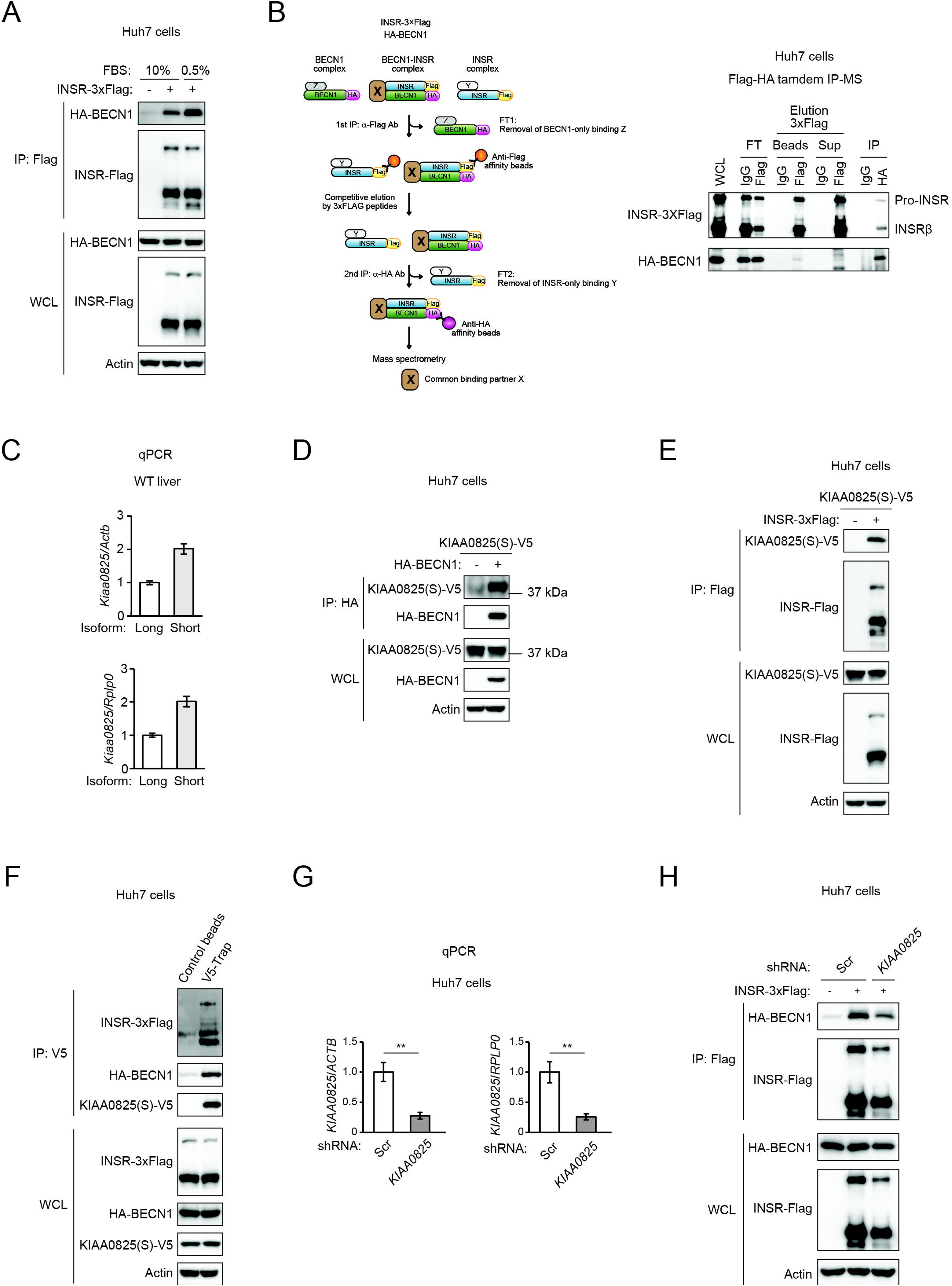
KIAA0825 is a binding partner of INSR and BECN1, and regulates INSR-BECN1-KIAA0825 complex formation. **(A)** Serum deprivation enhances BECN1-INSR interaction. Co-IP of BECN1 with INSR in Huh7 cells cultured with 10% or 0.5% FBS for 8 h. **(B)** An INSR-Flag HA-BECN1 tandem IP system in Huh7 cells to identify new complex components that simultaneous bind both BECN1 and INSR via mass spectrometry. The tandem IP allows for the removal of proteins that bind to BECN1 only, bind to INSR only, and bind to beads non-specifically. FT, flow-through; beads, beads-precipitated proteins; sup, protein pulldown dissolved in supernatant after first IP (Flag); IP, final precipitation after second IP (HA). **(C)** qPCR analysis of long and short isoforms of KIAA0825 in the liver of RD-fed WT mice. N=8 mice. **(D)** Co-IP of KIAA0825(S) (short isoform) by BECN1 in Huh7 cells transfected with HA-BECN1 and KIAA0825(S)-V5. **(E)** Co-IP of KIAA0825(S) by INSR in Huh7 cells transfected with INSR-Flag and KIAA0825(S)-V5. **(F)** Co-IP of INSR and BECN1 by KIAA0825(S) in Huh7 cells transfected with INSR-Flag, HA-BECN1 and KIAA0825(S)-V5. **(G)** qPCR analysis of KIAA0825 knockdown efficiency in Huh7 cells. N=4-6. T-test. **, P<0.01. **(H)** Co-IP of HA-BECN1 by INSR-Flag in Huh7 cells stably expressing scrambled (Scr) or KIAA0825 shRNA. Empty vector (without INSR-Flag) transfection serves as a negative control.

**Extended Data Figure 6.**
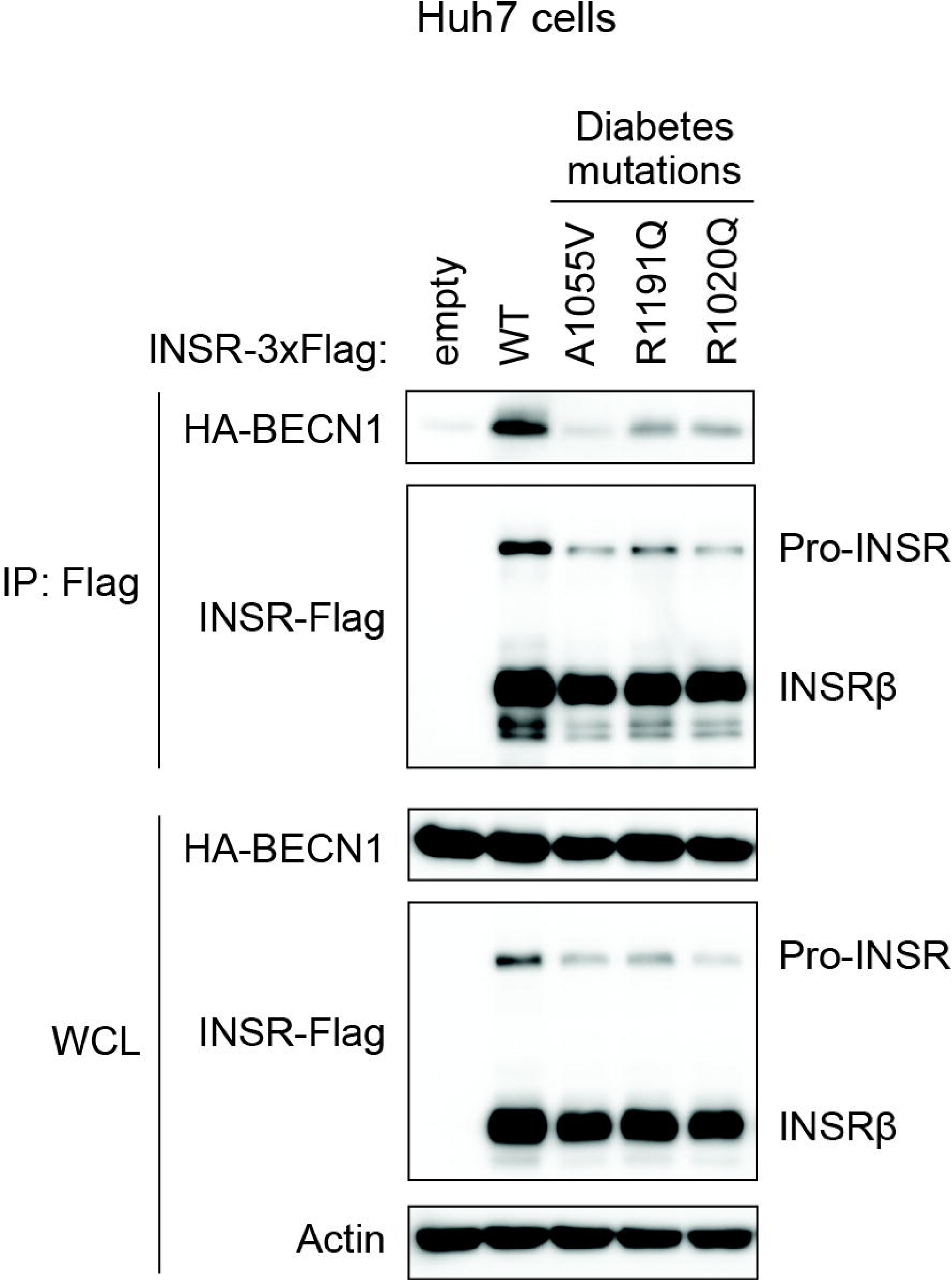
Monogenic diabetic INSR mutations impair binding of INSR to BECN1. Co-IP of BECN1 with human WT INSR or INSR mutants (A1055V, R1191Q and R1020Q identified in monogenic diabetic patients) in Huh7 cells overexpressing INSR-Flag and HA-BECN1.

## Discussion

In this study, we discovered novel regulators of INSR sorting, stability and signaling, and a new mechanism of INSR regulation by BECN1 and KIAA0825 (**Fig. 6E**). INSR mutations are single-gene defects (monogenic) causing childhood diabetes, but how INSR mutants lead to insulin resistance is poorly explored. Many INSR mutations causing monogenic diabetes locate in the intracellular kinase domain of subunit β, including R1020Q, A1055V and R1191Q that were studied here. Our data suggest that instead of affecting pro-INSR expression or directly impairing INSR kinase activity, these mutations impair insulin signaling by diverting mature INSR from cell-surface transport to EVs for secretion. The mutations block the key interaction of INSR with BECN1 and its binding partner KIAA0825. Given the current lack of effective therapies for patients with monogenic diabetes, our findings advance the mechanistic understanding of the disorder, and indicate that restoration of the INSR-BECN1-KIAA0825 complex may rescue the cell-surface trafficking and stability of mutant INSR by preventing its recruitment into and secretion via EVs.

We found that under fed conditions, deleting BECN1 specifically in the liver is sufficient to cause systemic hyperglycemia as in monogenic diabetes, supporting the link between BECN1 and glucose metabolic regulation. BECN1 has been reported to function in autophagic degradation and endocytic trafficking pathways ^28^. In this study, we revealed a new function of BECN1 in metabolism by preventing MVB sorting and EV secretion of INSR, different from its typical degradative roles in autophagy or endocytic trafficking. Our proteomic study unveiled that BECN1 carries out this function by forming a complex with KIAA0825, encoded by a diabetes-risk gene of unknown function identified in GWASs in multiple human populations, including Middle Eastern and Chinese populations ^23, 24^. Notably, KIAA0825 is also previously reported as a risk gene causing autosomal recessive postaxial polydactyly ^19–22^, suggesting its role in early limb development and patterning. We now demonstrated a new role of KIAA0825 in the regulation of INSR cell-surface trafficking and availability in synergy with BECN1. Given the key developmental role of KIAA0825 in polydactyly, it is possible that the BECN1-KIAA0825 complex is also important for insulin signaling and metabolism during embryonic or postnatal development.

In the BECN1^Δliver^ mice, our findings suggest that INSR-containing EVs are absorbed by hepatic vascular ECs, leading to reduced cell-surface INSR on hepatocytes and increased INSR on vascular ECs in the liver, but not on vascular ECs in more distant organs. It is likely mediated by fast absorption of ECs during circulation or by a paracrine manner. Yet, the physiological consequence of accumulated INSR on hepatic vascular ECs remains an open question. In addition, besides the liver, whether INSR is regulated via the similar INSR-BECN1-KIAA0825 mechanism in other metabolic organs needs further investigation, because INSR sorting and transport in tissues such as muscle and adipose tissue also plays an important role in controlling systemic glucose levels. Using additional tissue-specific mouse models will provide more insights to the understanding of cell-specific and systemic glucose metabolism in monogenic diabetes.

## Acknowledgements

We thank the following facilities at Northwestern University Feinberg School of Medicine for technical support: the Proteomics Core for proteomics studies, the NUSeq Core for sequencing, and the Metabolomics Core Facility for metabolomics analysis of glucose uptake. The study is supported by NIH R01 DK113170, R01 DK123447, and R01 DA056720 (to C. H.).

## Competing Interest Declaration

The authors declare no competing interests.

### Methods Mice

All mouse care and procedures were performed in accordance with animal experimental protocols approved by the Northwestern University Institutional Animal Care and Use Committee (IACUC). All mice were housed on a 14-h/10-h light/dark cycle with ad libitum access to chow diet and water. C57BL/6J (JAX #000664) and Becn1^flox^ (JAX #028794) mice were acquired from The Jackson Laboratory. All experiments were performed using sex- and age-matched mice. Diet-induced obese mice were generated by feeding a high-fat diet (D12492, Research Diets) for 8 weeks. Regular diet–fed mice were analyzed at 10–14 weeks of age, and high-fat diet–fed mice were analyzed at 18 weeks of age. For the fasting experiment, mice were fasted for 6 or 24 hours with free access to water.

Blood was collected from the tail vein, and blood glucose levels were measured using a glucose meter (Contour Next EZ; Ascensia Diabetes Care). For in vivo insulin stimulation to examine INSR and Akt phosphorylation levels, mice were intraperitoneally injected with insulin (1.5 U/kg) after 4 hours of fasting, and tissues were collected 15 minutes after insulin injection.

### Adeno-associated virus (AAV) tail vein intravenous injection

AAV2/8-TBG-PI-Cre-rBG (#107787-AAV8) and AAV2/8-TBG-PI-Null-bGH (#105536-AAV8) viruses were purchased from Addgene. Viral aliquots were stored at −80 °C until tail vein intravenous (i.v.) injection. Seven-week-old mice were intravenously injected with AAV at a dose of 1 × 10¹¹ viral genomes (VG) per mouse.

### Treadmill exercise

Three weeks after AAV injection, exercise was performed using a 10° uphill Exer 3/6 open treadmill (Columbus Instruments). Acute exercise (total 90 min) was performed as previously described ^8, 29^. Briefly, for acclimation, on day 1, mice ran for 5 min at 8 m/min, and on day 2, mice ran for 5 min at 8 m/min followed by 5 min at 10 m/min. On day 3, mice were subjected to acute running exercise starting at 12 m/min for 40 min. After 40 min, treadmill speed was increased by 1 m/min every 10 min for 30 min, followed by increases of 1 m/min every 5 min for 20 min.

Maximal running distance was measured following the acute exercise protocol described above with minor modifications. After the 90-min acute exercise, treadmill speed was increased by 1 m/min every 5 min until mice reached exhaustion.

### Measurement of glycogen content

Liver glycogen content was measured as previously described ^30, 31^. Briefly, 50–90 mg of mouse liver tissue was homogenized in 0.03 N HCl at 20 μl per mg of tissue. Homogenates (200 μl) were incubated with 200 μl of 2 M HCl for 2 hours at 95 °C and then neutralized by adding 200 μl of 2 M NaOH with vortexing. After centrifugation at 20,000 × g for 10 min at 4 °C, 20 μl of 1 M Tris-HCl (pH 7.4) was added, and glucose concentration was measured as total glucose.

To measure free glucose, 200 μl of homogenate was immediately neutralized with 2 M NaOH after adding 200 μl of 2 M HCl, followed by centrifugation at 20,000 × g for 10 min at 4 °C. After adding 20 μl of 1 M Tris-HCl (pH 7.4), glucose concentration was measured. Liver glycogen content was calculated by subtracting free glucose from total glucose.

### Total lipid extraction and triglyceride (TG) measurement

Total lipids were extracted from mouse liver using the Bligh and Dyer method as previously described ^8^. Extracted lipids were resuspended in isopropanol. Triglyceride (TG) levels were measured using Infinity Triglycerides Liquid Stable Reagent (TR22421, Thermo Fisher Scientific) according to the manufacturer’s instructions.

### In vivo 2-Deoxy-D-glucose (2-DG) uptake measurement

Mice were fasted for 6 hours (8 a.m. to 2 p.m.) and intraperitoneally injected with insulin (0.75 U/kg), 2-DG (0.15 g/kg), and D-glucose (1.35 g/kg). Thirty minutes after injection, mice were euthanized and perfused with PBS. Tissues were harvested and snap-frozen within 5 min. Metabolites were extracted by homogenizing in −80 °C–cooled 80% (vol/vol) methanol at 20 μl/mg of tissue. The homogenates were diluted 1:4 with −80 °C–cooled 80% (vol/vol) methanol and incubated at −20 °C overnight, followed by centrifugation at 20,000 × g for 15 min at 4 °C. Metabolite supernatants were analyzed at the Metabolomics Core Facility of Northwestern University.

### Plasmids

Double-stranded oligos encoding shRNAs against target genes were cloned into the AgeI and EcoRI restriction sites of the pLKO.1-TRC plasmid (Addgene plasmid #10878) or Tet-pLKO-puro (Addgene plasmid #21915). The sequences of gene-specific shRNAs used in this study were as follows: human BECN1 #1, CCCGTGGAATGGAATGAGATT; human BECN1 #2, CTCAAGTTCATGCTGACGAAT; human KIAA0825, CAACCCTAAGTGGAACATCTT; human RAB27A, CGGATCAGTTAAGTGAAGAAA. Scramble shRNAs in pLKO.1 and Tet-pLKO-puro (Addgene plasmids #1864 and #47541) were used as negative controls.

For CRISPR/Cas9 knockout plasmids, double-stranded oligos encoding gRNAs against target genes were cloned into the BbsI sites of the pSpCas9(BB)-2A-Puro (PX459) V2.0 plasmid (Addgene plasmid #62988). Gene-specific gRNA sequences were as follows: human BECN1, GGACACGAGTTTCAAGATCC; human KIAA0825, AATGAAGAGACGTCCTAGAG.

TagRFP-T-hBECN1, hBECN1-Tag-T-RFP, and HA-BECN1 were subcloned into the EcoRI site. INSR-3xFlag WT and patient-derived mutants were subcloned into the NheI site of pcDNA3.1(−) using the In-Fusion cloning kit (6389110, Takara Bio). hINSR-GFP was subcloned into the NheI–NotI site of pBRPB CAG-mCherry-IH using the In-Fusion cloning kit. pBRPB CAG-mCherry-IH was generated by replacing the puromycin resistance gene of pBRPB CAG-mCherry-IP (Addgene #106333) with the hygromycin resistance gene. pKT2/CAGXSP/CD63mScarlet (#182972) and hINSR-GFP (#22286) were purchased from Addgene.

For KIAA0825 gene cloning, cDNA libraries were constructed from total RNA of HEK293T and WI-38 cells using the SuperScript IV First-Strand Synthesis System kit (18091050, Thermo Fisher Scientific). Both C-terminal V5-tagged human KIAA0825 long and short isoforms were cloned from the cDNA libraries using the following primers: KIAA0825-V5-5’ primer, TGCTGGATATCTGCAGAATTCGCCACCATGGATTGGGATGATGAATATTCTCATAATTC;

KIAA0825-V5 long-isoform-3’ primer, TAGTCCAGTGTGGTGGAATTCTTACGTAGAATCGAGACCGAGGAGAGGGTTAGGGATAGGCTTACCCATGGTGGCGGATCCGCCGCCGCCCTGTTCCTCTATGTTATCTGAGGCAGAAG;

KIAA0825-V5 short-isoform-3’ primer, TAGTCCAGTGTGGTGGAATTCTTACGTAGAATCGAGACCGAGGAGAGGGTTAGGGATAGGCTTACCCATGGTGGCGGATCCGCCGCCGCCACCCAAAGCATGCACTGCTCCTCTATGC.

### Cells

Huh7, HepG2, HEK293T, WI-38 cells, and C2C12 myoblasts were cultured in Dulbecco’s modified Eagle’s medium (DMEM) with 10% fetal bovine serum (FBS), 100 units/ml penicillin, and 100 μg/ml streptomycin. For differentiation into myotubes, fully confluent C2C12 myoblasts (day 0) were cultured in DMEM containing 2% horse serum, 100 units/ml penicillin, and 100 μg/ml streptomycin, with medium refreshed every other day. On day 6, cells were used for experiments.

For tetracycline-inducible shRNA knockdown, cells were treated with 1 μg/ml tetracycline for 5 days before use. INSR-GFP–stably expressing HepG2 cells were generated using the PiggyBac transposon system. pBRPB CAG-INSR-GFP-IH and mouse codon-optimized PiggyBac transposase were transfected into HepG2 cells using Lipofectamine 3000, followed by selection with 400 μg/ml hygromycin. Cells with similar GFP brightness were collected by flow cytometry. Gene-specific knockout cells were generated by transfecting CRISPR/Cas9 knockout plasmids, followed by puromycin selection.

For insulin stimulation, cells were incubated in DMEM containing 0.5% FBS for 6 hours, then 100 nM insulin was added for the indicated time. After stimulation, cells were washed with PBS and lysed in Laemmli buffer.

### Lentivirus production and infection

HEK293T cells were transfected with a lentiviral vector together with pCMV-VSV-G (Addgene plasmid #8454) and psPAX2 (Addgene plasmid #12260) using Lipofectamine 3000 (L3000, Thermo Fisher Scientific). After 14–16 hours, the medium was replaced with fresh medium. Forty-eight hours post-transfection, the medium containing lentivirus was collected and filtered. Target cells were infected by incubation in lentivirus-containing medium with 8 μg/ml polybrene, followed by selection with appropriate antibiotics.

### Extracellular vesicle (EV)/exosome isolation

EVs were isolated from 20 ml of conditioned medium from Huh7 and HepG2 cells, and from 200 μl of pooled mouse plasma from four mice, as previously described ^32^. Briefly, cells were cultured in 150-mm dishes for 48 hours in 20 ml DMEM containing 10% exosome-free FBS. Mouse plasma was collected using EDTA-treated syringes. Conditioned medium and plasma were sequentially centrifuged at 300 × g for 5 min, then 3,000 × g for 10 min. Supernatants were centrifuged at 10,000 × g for 30 min at 4 °C, followed by 100,000 × g for 75 min at 4 °C using an ultracentrifuge (Optima XPN-100, Beckman Coulter). Pelleted vesicles were washed by resuspending in PBS and centrifuged again at 100,000 × g for 75 min at 4 °C. EVs were resuspended in 50 μl PBS for cell treatment or 60 μl Laemmli buffer for immunoblotting.

For EV treatment, differentiated C2C12 myotubes in 48-well plates were incubated with EV-containing medium (15 μl EV suspension in 1% FBS/DMEM) for 16 hours. Exosome-free FBS was prepared by ultracentrifugation at 100,000 × g for 75 min at 4 °C, followed by filtration through a 0.22-μm filter.

### Quantitative PCR (qPCR)

Total RNA was isolated from mouse liver and Huh7 cells using TRIzol Reagent (15596018, Thermo Fisher Scientific), and 2 μg of total RNA was reverse-transcribed using the High-Capacity cDNA Reverse Transcription Kit (4368814, Thermo Fisher Scientific) according to the manufacturer’s instructions. qPCR was performed using PowerUP SYBR Green Master Mix on a QuantStudio 6 Pro Real-Time PCR System (Thermo Fisher Scientific). Relative gene expression was calculated using the 2^-ΔΔCt^ method. The following gene-specific primers were used; mouse *Insr*-5’ TGCAAACAGATGCCACTAATCC; mouse *Insr*-3’ CAGACTCAAAGGGTGGGGAC; mouse *KIAA0825* (*2210408I21Rik*)-5 long isoform-5’ TCAGCCTCCTGCTTTTAACCC; mouse *KIAA0825* (*2210408I21Rik*)-5 long isoform-3’ ATCTTTCCTCATTTCCCACCTGT; mouse *KIAA0825* (*2210408I21Rik*)-5 short isoform-5’ ACAAGCAAGAGCTGCAACAAG; mouse *KIAA0825* (*2210408I21Rik*)-5 short isoform-3’ AGAGTTGATTTACGCTCTGGT; mouse *Rplp0*-5’ TTCGTGTTCACCAAGGAGGAC; mouse *Rplp0*-3’ ATGATCAGCCCGAAGGAGAAG; mouse *Actb*-5’ GATCTGGCACCACACCTTCT; mouse *Actb*-3’ GGGGTGTTGAAGGTCTCAAA; human *INSR*-5’ GGACCAGGCATCCTGTGAAAAT; human *INSR*-3’ TCAGGGGTGGGTCAATGTCT; human *KIAA0825*-5’ TGAAATAAACAATTCACAGCAAAAA; human *KIAA0825*-3’ TTCACCATTGAAGATGGAGCTA; human *GAPDH*-5’ ACCATCTTCCAGGAGCGAGAT; human *GAPDH*-3’ ATGACGAACATGGGGGCATC; human *ACTB*-5’ ACAGAGCCTCGCCTTTGC; human *ACTB*-3’ CCACCATCACGCCCTGG; human *RPLP0*-5’ GTGTTCGACAATGGCAGCAT; human *RPLP0*-3’ CAAGAAGGCCTTGACCTTTTCA.

### Immunoblotting

Mouse tissues or PBS-washed cultured cells were lysed in RIPA buffer (20 mM Tris-HCl [pH 7.4], 150 mM NaCl, 1% NP-40, 0.1% SDS, 0.5% sodium deoxycholate, 1 mM EDTA, 5 mM sodium fluoride, 1 mM sodium orthovanadate, 1 mM sodium pyrophosphate, 1 mM β-glycerophosphate, 1× protease inhibitor cocktail [P8340, MilliporeSigma]). Lysates were centrifuged at 15,000 × g for 10 min at 4 °C, and supernatants were collected. Protein concentrations were determined using a BCA assay kit (23225, Thermo Fisher Scientific). Proteins were denatured in Laemmli buffer by heating at 95 °C for 5 min. Equivalent amounts of protein were resolved by SDS-PAGE and transferred to PVDF or nitrocellulose membranes. Membranes were blocked with 0.5% skim milk or 5% BSA in TBS-T (Tris-buffered saline with 0.05% Tween-20) and probed with primary antibodies overnight at 4 °C. After washing three times for 10 min each with TBS-T, membranes were incubated with HRP-conjugated secondary antibodies for 2 hours at 4 °C. Specific bands were visualized by chemiluminescence using the ChemiDoc MP Imaging System (Bio-Rad), and band intensities were quantified using ImageJ.

### Co-immunoprecipitation (Co-IP)

Mouse liver was lysed in Triton X-100 lysis buffer (1% Triton X-100, 20 mM HEPES [pH 7.4], 150 mM NaCl, 1 mM EDTA, 5 mM sodium fluoride, 1 mM sodium orthovanadate, 1 mM sodium pyrophosphate, 1 mM β-glycerophosphate, 1× protease inhibitor cocktail [P8340, MilliporeSigma]). Lysates were centrifuged at 15,000 × g for 10 min at 4 °C, and supernatants were pre-cleared with 3 μg mouse IgG isotype control (10400C, Thermo Fisher Scientific) and Protein A/G beads (sc-2003, Santa Cruz Biotechnology) for 1 hour at 4 °C. 6 mg of total protein were incubated with 4 μg anti-INSR/Insulin Receptor β antibody (sc-57342, Santa Cruz Biotechnology) or mouse IgG control and Protein A/G beads for 14–16 hours at 4 °C. Beads were washed five times with lysis buffer, and immunoprecipitated proteins were eluted in Laemmli buffer for immunoblotting.

Huh7 cells transfected with plasmids using Lipofectamine 3000 were lysed in the same Triton X-100 buffer. Lysates were centrifuged at 15,000 × g for 10 min at 4 °C and incubated with target-specific antibodies and Protein A/G beads (sc-2003), anti-Flag M2 affinity gel (A2220, Millipore Sigma), or V5-Trap magnetic agarose beads (v5tma, Proteintech) for 14–16 hours at 4 °C. Beads were washed five times with lysis buffer, and immunoprecipitated proteins were eluted in Laemmli buffer and analyzed by immunoblotting.

Membrane fractions were prepared by centrifugation at 20,000 × g for 20 min at 4 °C following lysis in 0.25 M sucrose buffer (0.25 M sucrose, 1 mM EDTA, 20 mM HEPES [pH 7.4], 5 mM sodium fluoride, 1 mM sodium orthovanadate, 1 mM sodium pyrophosphate, 1 mM β-glycerophosphate, protease inhibitor cocktail) using a 27G needle.

For tandem immunoprecipitation, lysates were first incubated with anti-FLAG or IgG control beads for 2 hours at 4 °C, eluted with 3× FLAG peptide, and then incubated with anti-HA antibody or IgG control and Protein A/G beads overnight at 4 °C. Immunoprecipitated proteins were eluted in Laemmli buffer and analyzed by proteomics.

### Immunofluorescence staining

Livers and brown adipose tissue (BAT) were harvested from mice perfused with 4% paraformaldehyde (PFA) in PBS, cut into 5-mm-thick pieces, and fixed in 4% PFA/PBS for 4 hours. Tissues were incubated sequentially in 15% and 30% sucrose/PBS overnight for cryoprotection, embedded in optimal cutting temperature (OCT) compound, and sectioned at 15 μm using a cryomicrotome (CM1860, Leica Biosystems). Sections were washed twice with PBS for 5 min and once with PHEM buffer (60 mM PIPES, 25 mM HEPES, 10 mM EGTA, 2 mM MgCl_₂_, pH 6.9) for 5 min, followed by blocking in 5% normal goat serum (NGS) with 0.3% Triton X-100 in PHEM buffer for 60 min at room temperature. Sections were incubated with primary antibodies in antibody buffer (1% BSA, 0.3% Triton X-100 in PHEM) overnight at 4 °C, washed three times with PHEM buffer, and incubated with Alexa Fluor-conjugated secondary antibodies for 2 hours at room temperature. Sections were mounted with ProLong Diamond Antifade mountant (P36961, Thermo Fisher Scientific).

Cultured cells were washed three times with PBS, fixed and permeabilized in methanol at −20 °C for 10 min, washed with PHEM buffer, and blocked for 60 min at room temperature. Cells were incubated with primary antibodies in antibody buffer overnight at 4 °C, washed three times with PHEM-T (0.05% Tween-20 in PHEM), incubated with Alexa Fluor-conjugated secondary antibodies for 2 hours at room temperature, and mounted with ProLong Diamond Antifade mountant. Fluorescence images were acquired using a spinning-disk confocal microscope (X-Light V2 spinning disk confocal system [CrestOptics] with Hamamatsu Flash 4 camera [Hamamatsu Photonics] mounted to a Nikon ECLIPSE Ti microscope [Nikon]).

### Antibodies

The following antibodies were used in immunoblotting, immunoprecipitation, and immunofluorescence: anti-Flag M2 antibody (F1804) (Millipore Sigma); anti-CD31 (553370) antibody (BD Biosciences); anti-CD63 (ab217345) antibody (Abcam); anti-Alix (12422-1-AP), anti-BECN1 (11306-1-AP), anti-CD9 (20597-1-AP), anti-Flotillin1 (15571-1-AP), anti-GLUT2 (20436-1-AP), anti-RFP (5f8), and anti-TSG101 (28283-1-AP) antibodies (Proteintech); anti-Alix (sc-53540), anti-BECN1 (sc-48341), anti-CD63 (sc-5275), anti-CRY1 (sc-393466), anti-GAPDH (sc-365062), anti-HA-tag (sc-7392), anti-Insulin R β (INSRβ) (sc-57342), anti-LAMP1 (sc-20011), anti-pan cadherin (sc-59876), anti-RAB27A (sc-74586), and HRP-conjugated β-actin (sc-4778 HRP) antibodies (Santa Cruz Biotechnology); anti-Akt (4691), anti-BiP (3177), anti-Calnexin (2679), anti-EGF Receptor (4267), anti-HA-tag (3724), anti-IGF-I Receptor β (9750), anti-Insulin receptor beta (INSRβ) (23413), anti-IRS1 (3407), anti-Phospho-IGF-I Receptor beta (Tyr1135)/Insulin Receptor beta (Tyr1150) (3918), anti-p70 S6 kinase (2708), anti-phospho-Akt (Ser473) (4060), anti-phospho-mTOR (Ser2448) (5536), anti-mTOR (2983), and anti-V5 (13202), anti-SQSTM1/p62 (5114) antibodies, and HRP-conjugated anti-Mouse IgG light-chain specific (91196) antibodies (Cell Signaling Technology); HRP-conjugated anti-Rabbit IgG (111-035-003), HRP-conjugated anti-Mouse IgG (115-035-003), and HRP-conjugated anti-Mouse IgG light chain specific (115-035-174) antibodies (Jackson ImmunoResearch); Alexa Fluor Plus 488-conjugated F(ab’)_₂_ anti-Mouse IgG (A48286), Alexa Fluor Plus 594-conjugated F(ab’)_₂_ anti-Rabbit IgG (A48284), and Alexa Fluor Plus 647-conjugated anti-Rat IgG (A48265) (Thermo Fisher Scientific).

### Quantification and Statistical Analysis

All data are presented as mean ± standard error of the mean (SEM). Statistical comparisons between two groups were performed using Student’s t-test. For comparisons involving more than two groups, two-way analysis of variance (ANOVA) followed by Tukey’s post hoc test was used. Analyses were conducted using GraphPad Prism (Dotmatics). Differences with a p-value < 0.05 were considered statistically significant.

